# A Pmk1-Regulated Mst12-Bip1 Network Coordinates Appressorium Function, Effector Deployment, and Invasive Growth by *Magnaporthe oryzae*

**DOI:** 10.64898/2026.09.11.750954

**Authors:** Camilla Molinari, Neha Sahu, Lauren Ryder, Xia Yan, Vangeli Geshkovski, Tatsuya Nobori, Nicholas J. Talbot

**Affiliations:** The Sainsbury Laboratory, Norwich Research Park, Colney Lane, Norwich, NR4 7UH, United Kingdom

**Keywords:** Plant pathogen, rice blast, appressorium, effectors, gene expression

## Abstract

To cause rice blast disease, the filamentous fungus *Magnaporthe oryzae* develops a specialised infection cell called an appressorium, which generates enormous turgor to breach the rice leaf cuticle. Although key regulators of appressorium development have been identified, it is not known how they drive the extensive transcriptional reprogramming required for infection-related morphogenesis. Here, we show that the Pmk1 MAP kinase orchestrates plant infection by regulating a transcriptional network controlled by the Mst12 and Bip1 transcription factors. Bip1 is regulated both by Pmk1-dependent phosphorylation and at the transcriptional level, while Mst12 binds to a *cis*-acting element upstream of *BIP1* essential for pathogenesis. Bip1 and Mst12 are both necessary for regulating a set of Pmk1-dependent appressorium-specific genes, including transcriptional regulators, cell wall-degrading enzymes, and effector proteins. In addition, Mst12 specifically regulates functions required for appressorium-mediated penetration, while Bip1 controls a distinct sub-set of effector genes deployed during invasion of plant tissue. When considered together, these findings define a Pmk1-dependent transcriptional network required for plant infection by the blast fungus.

## Introduction

Appressorium-mediated infection is a key innovation used by many of the world’s most devastating plant pathogenic fungi to infect intact host plants (Talbot, 2019, Ryder *et al*., 2022). In *M. oryzae* appressorium development is a complex process accompanied by enormous transcriptional reprogramming, involving almost 50% of the genome being differentially expressed (Osés-Ruiz *et al*., 2021, Sakulkoo *et al*., 2018), yet how the regulatory pathways controlling this process operate remains poorly understood.

During *M. oryzae* infection, a germinating conidium differentiates a dome-shaped appressorium that generates enormous turgor pressure, enabling formation of a penetration peg to mechanically rupture the leaf cuticle (Ryder et al., 2022, Wilson & Talbot, 2009). Once inside the plant, *M. oryzae* colonises epidermal cells through formation of bulbous invasive hyphae and deployment of a repertoire of secreted effector proteins that manipulate plant physiology and suppress immunity (Oliveira-Garcia *et al*., 2024). *M. oryzae* is predicted to encode at least 546 effectors, predominantly expressed between 24h-48h post-infection (Yan *et al*., 2023). Infection-related morphogenesis and effector deployment are coordinated by interconnected signalling pathways which are subject to cell cycle regulation, and control cytoskeletal re-modelling, autophagy-associated cell death of the conidium, polar/non-polar growth transitions, and cell wall biogenesis (Wang *et al*., 2024, Ryder et al., 2022, Wengler & Talbot, 2025, Wilson & Talbot, 2009).

In *M. oryzae,* appressorium formation, host penetration, and plant tissue invasion are all regulated by the Pmk1 MAPK signalling pathway, which is conserved among diverse plant-pathogenic fungi irrespective of whether they form an appressorium to infect their host (Xu & Hamer, 1996, Turrà *et al*., 2014, Sakulkoo et al., 2018, Cruz-Mireles *et al*., 2021). Pmk1 is activated when *M. oryzae* senses a hard hydrophobic surface and perceives plant-derived signals via extracellular sensors Msb2 and Sho1 and associated Ras GTPases (Liu *et al*., 2011). This triggers a three-tiered MAPK module in which the Mst11 MAPKKK activates the Mst7 MAPKK, which in turn phosphorylates Pmk1 to initiate appressorium development. Without Pmk1, mutants are unable to form an appressorium (Xu & Hamer, 1996), while conditional inactivation of Pmk1, by generation of an analogue-sensitive *pmk1^AS^* allele, impairs the ability of *M. oryzae* to move from cell-to-cell inside plant tissue (Sakulkoo et al., 2018).

Comparative transcriptional profile analysis has shown that 6333 genes (∼ 49% of the *M. oryzae* genome) are differentially regulated in a *Δpmk1* mutant compared to an isogenic wild type strain, a large proportion of which are also expressed specifically during appressorium development (Osés-Ruiz et al., 2021). Phosphoproteomic analysis furthermore revealed 2062 phosphoproteins associated with appressorium development, many of which are Pmk1-dependent or direct substrates of the kinase (Cruz-Mireles *et al*., 2024). Among identified Pmk1 substrates are transcriptional regulators such as Mst12 and Hox7 that are required for pathogenicity (Cruz-Mireles et al., 2024, Osés-Ruiz et al., 2021). This suggests that Pmk1 regulates a hierarchy of transcription factors that control infection related morphogenesis, but how these regulators and their targets orchestrate appressorium development and invasive growth is not known.

In this study we set out to investigate pathogenicity-related transcriptional regulation, focusing on the connection between regulators of morphogenetic development and virulence. We report that Bip1 (<u>B</u>-ZIP Involved in <u>P</u>athogenesis-1), a recently described transcription factor in *M. oryzae* (Lambou *et al*., 2024) is controlled by the Pmk1 MAPK signalling pathway and Mst12, forming a transcriptional network essential for blast disease. Previously, it was reported that *Δbip1* mutants can form appressoria but are unable to penetrate host tissue, suggesting a function in plant tissue colonisation (Lambou et al., 2024). It was also proposed that Bip1 functions independently of both Pmk1 and Mst12 leaving its precise regulatory role unknown (Lambou et al., 2024). By contrast, we present evidence that Bip1 is regulated by Pmk1-dependent phosphorylation, which is essential for its role in blast disease. Structural modelling and *in vitro* binding studies also provide evidence that Mst12 regulates *BIP1* expression and that this too is critical for fungal pathogenesis. Global transcriptional profiling and network analysis reveal that Bip1 and Mst12 in fact regulate a large overlapping set of genes during plant infection, but Bip1 additionally regulates specific functions associated with plant tissue colonisation, including a set of effector-encoding genes. When considered together, this study offers new insight into how a Pmk1-Mst12-Bip1 regulatory network coordinates the morphogenetic and physiological transitions required for blast disease.

## Results

### Bip1 is a transcription factor associated with infection-related development

We set out to explore the relationship between the Pmk1 MAP kinase and the Bip1 transcriptional regulator (Lambou et al., 2024). We first generated two independent *BIP1* deletion mutants in the Guy11 background, confirmed by whole-genome sequencing (Fig. S1). Both *Δbip1* mutants formed appressoria on hydrophobic surfaces but failed to penetrate host tissue, consistent with the previously reported phenotype (Lambou et al., 2024), and confirming that Bip1 is required for virulence by the blast fungus (Fig.1*A* and *B*). The *Δbip1* mutants also showed growth and sporulation defects and produced unusually white colonies (Fig. S2*A* and *B*), suggesting a possible defect in melanin biosynthesis reminiscent of *Δpmk1* and *Δmst12* mutants (Zhu *et al*., 2021). A Bip1– mNeonGreen gene fusion localised predominantly to nuclei during appressorium development, Bip1 co-localise with a histone H1-TdTomato nuclear marker and shows strongest signal during conidial germination and in mature appressoria (Fig. S3*A* and *B* and Video 1). Because both *Δmst12* and *Δbip1* mutants form appressoria (Fig. 1*D*) but cannot penetrate host tissue, we examined the cytoskeletal re-modelling essential for penetration peg formation. Fluorescent markers for actin, Septin 5, and the septin kinase Chm1 were introduced into Guy11, *Δmst12* and *Δbip1* mutants, respectively (Fig. 1*C* and *D*). In *Δmst12* mutants, each cytoskeletal marker was mislocalised, as previously reported (Gupta *et al*., 2015, Osés-Ruiz et al., 2021), confirming that Mst12 is required to organise septin ring formation for polarised growth and peg formation. By contrast, in *Δbip1* mutants, actin and Chm1 localise to the appressorium base, whereas Sep5 assembles into a disc rather than the septin ring observed in Guy11. Taken together, this provides evidence of a temporal or structural defect in septin aggregation and organisation in *Δbip1* mutants rather than the more extensive cytoskeletal defects observed in *Δmst12* mutants.

**Figure 1.**
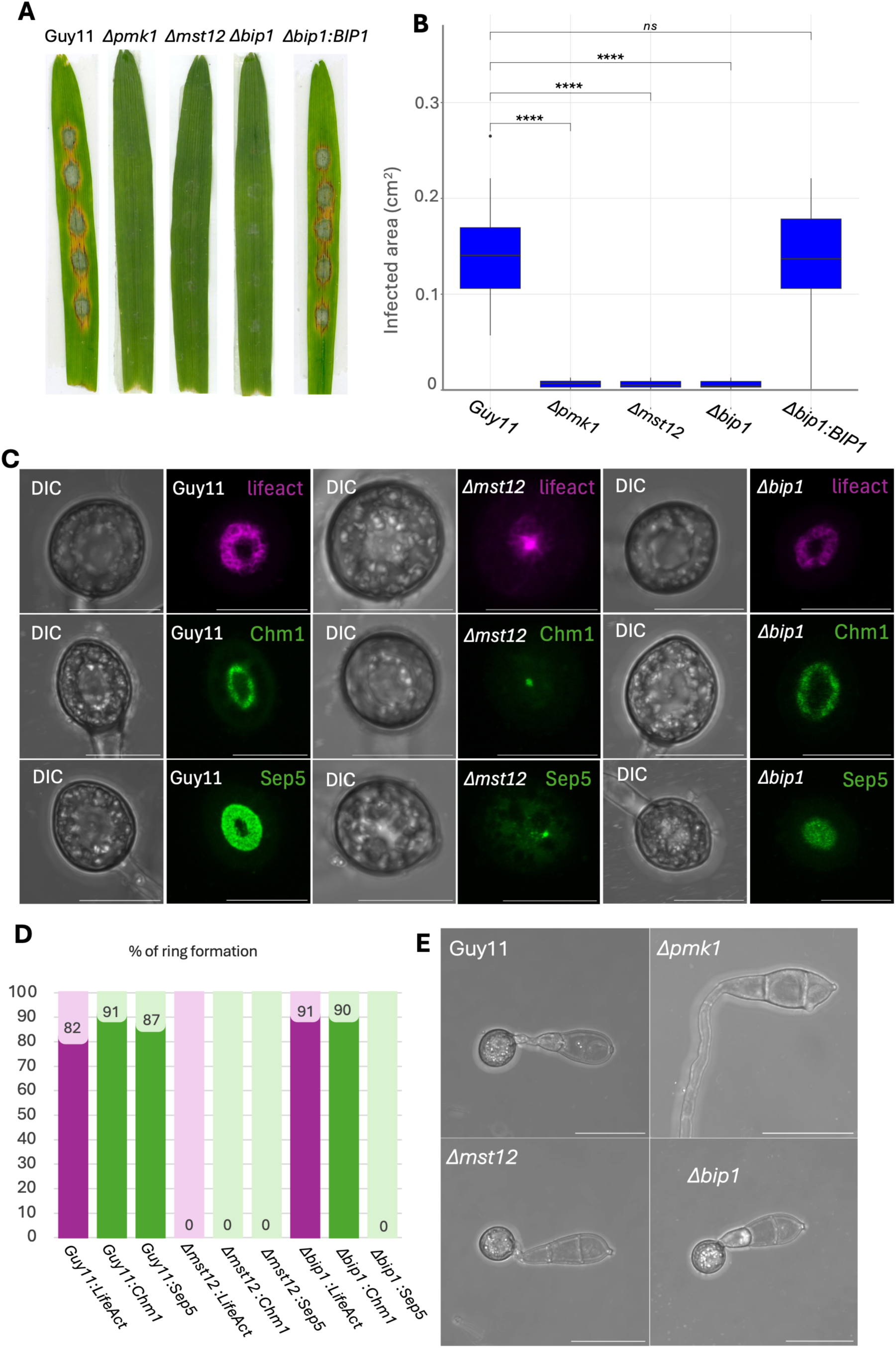
Bip1 is a transcription factor required for infection-related development. (A) Detached leaf assays (barley cv. Golden Promise; see Methods) comparing Guy11, *Δmst12, Δpmk1*, *Δbip1*, and *Δbip1*:*BIP1-GFP*, 5 dpi. (B) Box plots show the calculated disease area (n=20 lesions per replicate). The mean value is shown as a horizontal line inside the box. T-tests show significantly reduced virulence (p-value<0.05) for all mutant strains compared to wild-type infection. Results represent four biological replicates. (C) Actin (Lifeact-RFP), Chm1 (Chm1-GFP) and Sep5 (Sep5-GFP) form normal rings in Guy11 appressoria at 24 h but are extensively mislocalised in *Δmst12*; in *Δbip1*, actin and Chm1 localise normally but Sep5 forms a disc rather than a ring, quantified in panel D. Laser confocal microscopy at 24h, maximum Z-stack projection representative of three biological replicates. Scale bar = 10 μm. (E) Micrographs show representative appressorium morphologies for Guy11, *Δmst12, Δpmk1*, and *Δbip1 M. oryzae* strains. Conidia were harvested, inoculated on hydrophobic coverslips, and observed by laser confocal microscopy at 24h. Micrographs show a maximum projection of Z-stack images. Scale bar = 20 μm.

### Pmk1 is necessary for both transcriptional and post-translational regulation of Bip1

We observed that *BIP1* transcript levels were consistently reduced in a *Δpmk1* mutant relative to wild type across appressorium development (0–24 h; Fig. 2A), suggesting that *BIP1* expression is Pmk1-dependent at early stages of appressorium development (Osés-Ruiz et al., 2021). Examination of a time-resolved phosphoproteomic analysis of appressorium development of *M. oryzae,* comprising 2,062 phosphorylated proteins (Cruz-Mireles et al., 2024), furthermore identified a candidate MAPK phosphorylation site in Bip1 at serine 32, located in a poorly conserved region N-terminal to a predicted nuclear localisation signal, bZIP, and dimerisation domains (Fig. 2*B*). The corresponding phosphopeptide showed reduced abundance in a *Δpmk1* mutant compared to Guy11, particularly at later time points (4 and 6 h) during appressorium morphogenesis (Fig. 2*C*), consistent with Bip1 phosphorylation requiring Pmk1 activity, either directly or indirectly. We also observed that serine 32 is highly conserved in plant-associated fungi, in particular hemibiotrophic, necrotophic, and biotrophic fungal pathogens (Fig. 2*D*), suggesting a link between plant adaptation and phosphorylation of this Bip1 residue. Therefore to test the functional relevance of the phosphorylation event, we introduced *BIP1* alleles carrying amino acid substitutions that alter the phosphorylation status of the transcription factor, either through a non-phosphorylatable substitution (S32-A32) or a phosphomimetic (S32-D32) mutation, into a *Δbip1* mutant of *M. oryzae*. Transformants expressing non-phosphorylatable Bip1 were unable to restore the ability to cause blast disease of a *Δbip1* deletion mutant, whereas mutants expressing phosphomimetic Bip1 showed wild-type levels of blast disease (Fig. 2*E* and *F*). We conclude that Pmk1-dependent Bip1 phosphorylation is critical for *M. oryzae* host penetration and pathogenicity.

**Figure 2.**
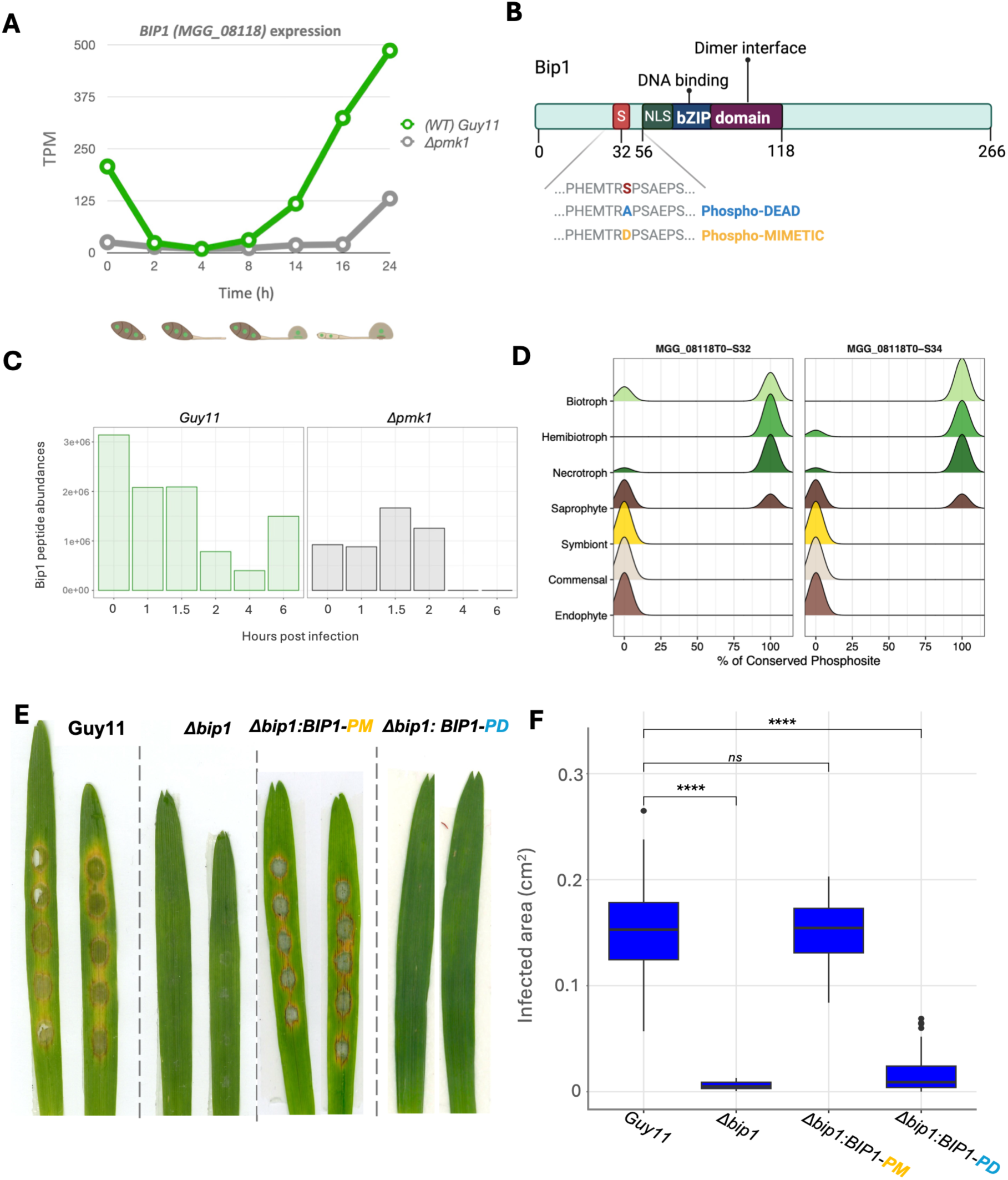
Bip1 is regulated by the Pmk1 pathway, transcriptionally and post translational by phosphorylation. (A) Cartoon of Bip1 protein with a phosphorylation site (serine in position 32) and bZIP domain from position 56 to 118, comprising an NLS, DNA-binding domain and dimer interfaces. Below is a representation of the chimeric phosphorylation sequences used to complement the *Δbip1* mutant. (Image created using Biorender.com). (B) *BIP1* transcript abundance (TPM) mean values across seven different time points (0, 2, 4, 8, 14, 16, and 24h) during appressorium development for two distinct strains: Guy11 (green) and *Δpmk1* mutant (grey) extracted from publicly available transcriptomic dataset (Osés-Ruiz et al., 2021). (C) Conservation of Bip1 serine 32/34 across 42 fungal species of differing lifestyles. (D) Relative phosphopeptide abundances for S32 of Bip1 across a time-course of appressorium development (0, 1, 1.5, 2, 4, and 6h) for Guy11 wild type and *Δpmk1* mutant. (E) Detached leaf assays (barley cv. Golden Promise; see Methods) comparing Guy11, *Δbip1, Δbip1:BIP1-PhosphoMIMETIC,* and *Δbip1:BIP1-phosphoDEAD*, 5 dpi. (B) Box plots show the calculated disease area (n=20 lesions per replicate). The mean value is shown as a horizontal line inside the box. T-tests show significantly reduced virulence (p-value<0.05) for *Δbip1 and Δbip1:BIP1-PhosphoDEAD* strains compared to wild-type infection. Results represent four biological replicates.

### An Mst12-interacting *BIP1* promoter element is required for fungal pathogenicity

As Bip1 appears to be regulated via the Pmk1 pathway, we next explored potential connections between Bip1 and the widely reported Pmk1 downstream target Mst12 (Park *et al*., 2002, Osés-Ruiz et al., 2021). Motif analysis of the *BIP1* promoter using Mst12 ChIP-seq binding data (Osés-Ruiz et al., 2021) identified a single Mst12-binding site, “CCTGTAAT,” 661 bp upstream of the *BIP1* start codon (Fig. 3C). AlphaFold3 protein–DNA modelling predicted that this motif is bound by the C_2_H_2_ zinc-finger region of Mst12, a domain absent from Mst12 orthologues in ascomycete yeasts, but reported to be involved in pathogenicity (Wong Sak Hoi & Dumas, 2010). To define the putative interaction in more detail, we modelled the C_2_H_2_ zinc finger domains with the *BIP1* promoter motif, which increased confidence scores compared to the full-length Mst12 protein, consistent with extensive disorder in other predicted regions of Mst12. Systematic modelling with truncated variants, for example, showed that the full-length protein yielded the highest interface score (0.9 ipTM) but low overall confidence (0.39 pTM), whereas restricting the model to the C_2_H_2_ zinc finger region maintained a high interface score (0.83 ipTM) but substantially improved the overall score (0.82 pTM). While models containing only the first or the second zinc finger reduced interface confidence (0.65-0.67 ipTM scores) (Fig. 3*B* and Fig. S4), supporting maximal predicted binding confidence is reached when both zinc fingers are present.

**Figure 3.**
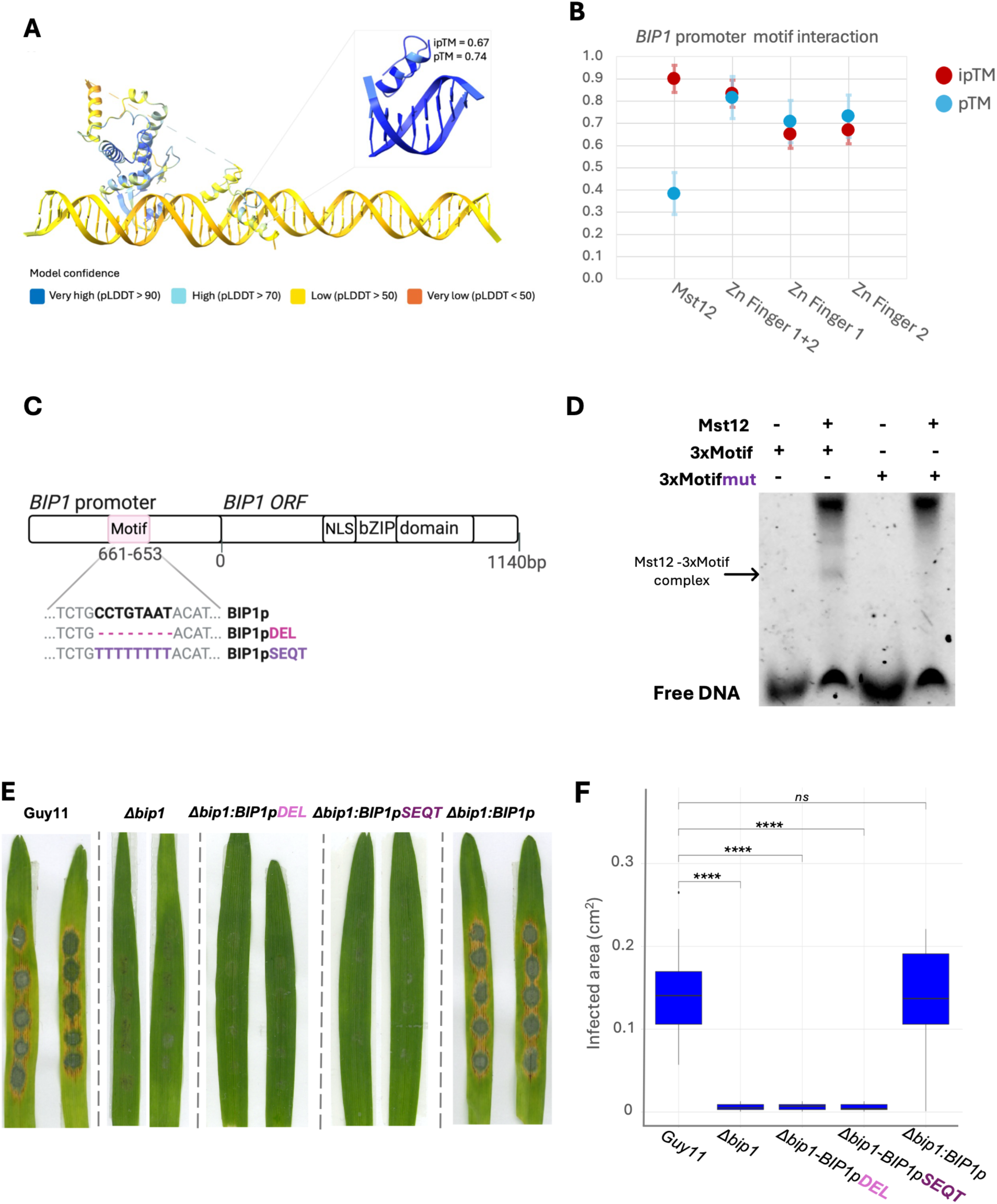
An Mst12-interacting *BIP1* promoter element is required for pathogenicity by the blast fungus. (A) Alphafold-3 prediction of Mst12 protein with 30bp of the *BIP1* promoter, containing a putative Mst12 DNA-binding motif (Osés-Ruiz et al., 2021). Contact is predicted to occur through the second C_2_H_2_ zinc finger domain of Mst12 and the predicted DNA-binding site on the *BIP1* promoter. pLDDT scores are shown with the model confidence colouring, and distorted regions have been cropped for visualisation purposes. (B) pTM and ipTM values for Alphafold-3 models of the *BIP1* promoter motif “CCTGTAAT” with full length Mst12, the C_2_H_2_ zinc finger region and each C_2_H_2_ zinc finger individually (5 seeds each) (C) Cartoon of *BIP1* promoter and coding region depicting the location of the Mst12 DNA-binding motif “CCTGTAAT” 661bp upstream of the start codon of *BIP1* with the promoter chimeras used to complement the *Δbip1* mutant. (Image created using Biorender.com). (D) EMSA: fluorescently labelled probes containing three tandem repeats of “CCTGTAAT” or a mutant “CAAATAAT” motif, incubated with recombinant Mst12 (1:2 or 1:10 molar ratio); binding is specific to the cognate motif and increases with molar ratio. (E) Detached leaf assays (barley cv. Golden Promise; see Methods) comparing Guy11, *Δbip1, Δbip1:BIP1pDEL, Δbip1: BIP1pSEQT,* and *Δbip1: BIP1p-BIP1-GFP*, 5 dpi. (F) Box plots show the calculated disease area (n=20 lesions per replicate). The mean value is shown as a horizontal line inside the box. T-tests show significantly reduced virulence (p-value<0.05) for all mutant strains compared to wild-type infection, except for the *Δbip1: BIP1p-BIP1-GFP* complementation. Results represent four biological replicates.

As an internal control, the experimentally validated interaction between Bip1 and its cognate DNA motif, “TGACTCG”, in the promoter of the *MGG_08381* gene (Lambou et al., 2024) was also modelled. The best-ranked Bip1–DNA motif complex showed an ipTM of 0.75 and a pTM of 0.32, lower than for the Mst12–*BIP1* motif complex. To probe specificity further, we generated additional models in which the *BIP1* motif was either deleted or replaced with a poly-thymine sequence and modelled an Mst12 splice variant lacking the second C_2_H_2_ domain. These predictions ranked lower than the wild-type configuration (Fig. S5 and Fig. S6), consistent with the second C_2_H_2_ zinc finger domain of Mst12 specifically recognising the ChIP-seq–defined “CCTGTAAT” motif found in the *BIP1* promoter.

To test the predicted binding of Mst12 to the “CCTGTAAT” motif we used an electrophoretic mobility shift assay (EMSA). Recombinant Mst12 was incubated at a 1:2 molar ratio with labelled oligonucleotides containing three tandem repeats of the *BIP1* promoter fragment or three tandem repeats of a mutant version of the DNA motif “CAAATAAT”. Mst12 showed binding to its cognate motif but not the mutant version (Fig. 3*D*) confirming formation of an Mst12-DNA complex. To test the biological relevance of this interaction, we constructed two *BIP1* promoter chimaeras: one in which the *BIP1* promoter motif was deleted and another in which the motif was replaced with a poly-T sequence. Each construct was used to complement the *Δbip1* mutant. In both cases, the tested chimeric promoter was unable to restore the ability of a *Δbip1* mutant to cause blast disease (Fig. 3*E* and *F*), while a non-chimeric version of the *BIP1* promoter restored pathogenicity. The Mst12 DNA-binding motif “CCTGTAAT” in the *BIP1* promoter is therefore required for penetration and blast disease. Through motif scanning, we identified a total of 658 genes containing this Mst12-binding motif in *M. oryzae*, including 22 further transcription factor-encoding genes, such as *BZIP11* and *MBF1* involved in virulence, and *BZIP2/GCN4*, reported to be essential for viability (Fan *et al*., 2017, Kong *et al*., 2015, Jeon *et al*., 2007, Tang *et al*., 2015), suggesting the Pmk1–Mst12 cascade may regulate a wider set of pathogenicity regulators through a specific promoter motif.

### Mst12 and Bip1 co-regulate a subset of genes during appressorium development

A previous microarray analysis suggested that Bip1 acts as a transcriptional activator during appressorium development and identified 40 putatively regulated genes (Lambou et al., 2024). To provide higher resolution analysis and determine how Bip1 acts as part of the Pmk1 MAPK signalling pathway, we performed global RNA-seq profiling of both *Δbip1* and *Δmst12* deletion mutants during appressorium development of *M. oryzae* (0, 4 and 24 h), identifying 4,791 and 5,437 differentially expressed genes (DEGs) respectively (log2 fold-change ≥ 1 and adjusted p-value ≤ 0.05 (Robinson *et al*., 2009, Pimentel *et al*., 2017, Bray *et al*., 2016)). We found 2,633 genes co-regulated Bip1 and Mst12, representing 55% of the Bip1 regulon and 48% of the Mst12 regulon, respectively (Fig. 4*A*). Both transcription factors acted as activators and repressors of gene expression. This is in marked contrast to the previous report in which Bip1 was only observed to be necessary for up-regulation of gene expression (Lambou et al., 2024). Mst12 was necessary for activation of 2,948 genes and repression of 2,489, with 335 genes showing biphasic (stage-dependent) behaviour; Bip1 was necessary for up-regulation of 2,896 genes and repression of 1,895, with 213 biphasic genes. Of Bip1-activated DEGs, the proportion containing the previously reported Bip1-binding motif “TGACTCG” (Lambou et al., 2024) in their promoters increased over developmental time (10% at 0 h, 11% at 4 h, 16% at 24 h), suggesting these represent direct Bip1 targets, while the remainder may be regulated indirectly or via additional Bip1-binding elements.

**Figure 4.**
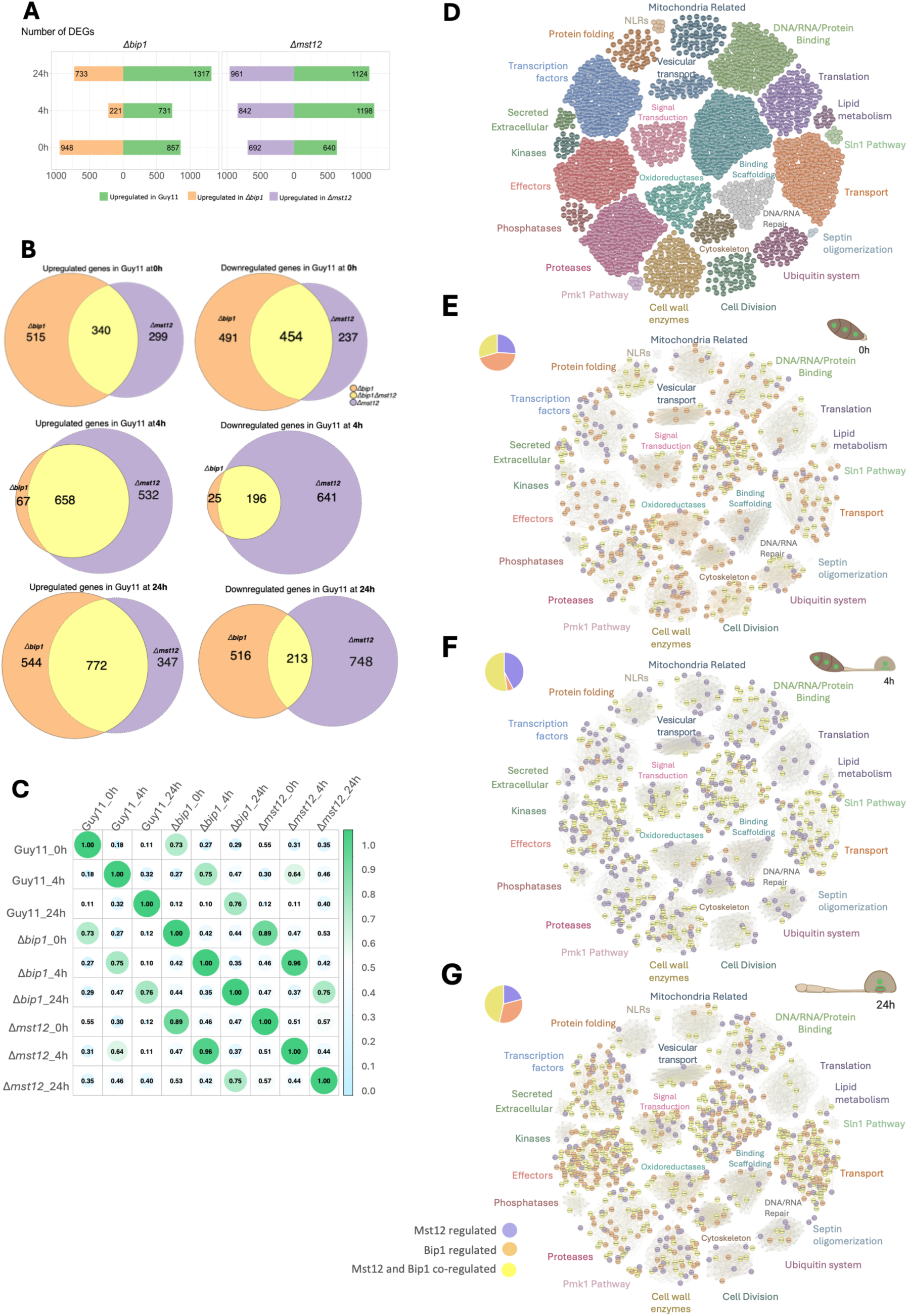
Bip1 and Mst12 show both co-regulatory and distinct gene regulation during appressorium development. (A) DEG counts across three time points (0, 4, 24 h) for *Δbip1* and *Δmst12*. (B) Euler plots of gene overlap across samples/time points *(Δbip1*; orange, *Δmst12*; purple, *Δbip1* and *Δmst12*; yellow). (C) Pearson correlation matrix of global transcriptomic profiles for Guy11, *Δmst12,* and *Δbip1* (green = high, light blue = low correlation). (D) Base functional network of DEGs, built from InterPro domains and manual annotation into 24 colour-coded clusters (full breakdown in SI Table 1), dominated by binding/scaffolding, transport, signal transduction, transcription factor, and effector-encoding genes. DEGs up-regulated in Guy11 relative to *Δbip1* (orange), *Δmst12* (purple), or both (yellow) are plotted onto this network at 0 h (E), 4 h (F), and 24 h (G).

Multidimensional scaling (MDS) explained 58% of the total variance across transcriptional fingerprints, with biological replicates clustering tightly (Fig. S7). Bip1 and Mst12 display a strong transcriptional overlap at all timepoints (Fig. 4*B* and Fig. S8) supported by a Pearson correlation matrix, though at 24h there is also a great divergence observed, when both factors co-regulate the largest subset of genes, but also display additional transcriptomic specificity (Fig. 4*C*). Biphasically co-regulated genes included known regulators of appressorium development and infection, such as the transcription factors *STU1*, and other bZIPs (Pham *et al*., 2015, Tang et al., 2015), the peroxisomal gene *HEX1* required for Woronin body septal pore function (Soundararajan *et al*., 2004), and the *MAS1*/*MAS3* genes necessary for biotrophic growth (Gong *et al*., 2022), among others (full list in SI Table 1). Individual biphasic regulation for Mst12 included a set of genes required for appressorium function and virulence such as developmental transcription factors, regulators of DHN-melanin biosynthesis and the NADPH oxidase complex (Lu *et al*., 2014, Tsuji *et al*., 2000, Deng *et al*., 2016, Wang *et al*., 2007, Galhano *et al*., 2017, Fu *et al*., 2014). While Bip1 stands out for regulating lipid biosynthesis genes such as *FAR1* (bin Yusof *et al*., 2014, Li *et al*., 2011), surface/environment sensing genes such as *PTH11* and related G-protein coupled receptors (DeZwaan *et al*., 1999, Ye *et al*., 2014). Collectively Mst12 and Bip1 control a wide range of cellular functions, that are essential for infection-related morphogenesis.

### Mst12 and Bip1 have co-regulatory and independent functions during appressorium-mediated plant infection

To identify the functional categories regulated by Mst12 and Bip1, we assigned DEGs to a network correlation map (Molloy *et al*., 2024) built from InterPro domains and manual annotation, yielding 24 functional clusters dominated by transcription factors, transporters, signal transduction components, cell wall enzymes, and 295 predicted effectors (Fig.4*D*). Overlaying Mst12-regulated (purple), Bip1-regulated (orange), and Mst12-Bip1-regulated (yellow) genes onto this network across the three developmental timepoints revealed a clear temporal pattern (Fig. *4E–G*, Fig. S9). In ungerminated conidia (0 h), Bip1 had a larger activating footprint than Mst12 (514 vs. 299 genes), dominated by primary metabolism and redox genes (e.g. *HYR1, BUF1*) (Que *et al*., 2020, Zhang *et al*., 2015, Huang *et al*., 2011, Chumley & Valent, 1990), while Mst12-activated genes were enriched for transcription factors (Cao *et al*., 2016, Lu et al., 2014). Both Bip1 and Mst12-regulated genes (340) included genes for autophagy-mediated recycling (*ATG6, ATG27*) (Wengler & Talbot, 2025) and stress responses (*SOD1, HEX1*) (Li *et al*., 2023, Soundararajan et al., 2004) essential for germination. From 4 h, as appressoria began to differentiate, Mst12 became the dominant regulator, activating 532 genes encoding core appressorium and penetration machinery (e.g. melanin biosynthesis, septin/cytoskeletal regulators, the Hox7 transcription factor) (Jacob *et al*., 2014, Takemoto *et al*., 2011, Osés-Ruiz et al., 2021, Peng *et al*., 2011), while the unique contribution of Bip1 at this stage was limited, but included velvet-complex members (*VEA, VOSA*) (Meng *et al*., 2023). The shared set of regulated genes (658) included secreted effector candidates, components of the Pmk1 signalling pathway (*RPP2, RPP3*), and melanin/polyketide synthases (Chen *et al*., 2022, Irieda *et al*., 2019, Zhang *et al*., 2024, Osés-Ruiz et al., 2021) further advancing germination and appressorium generation required for virulence.

By the time of appressorium maturation (24 h), the Bip1–Mst12 co-regulated set was largest (771 genes), enriched for effectors (60 genes), transporters (31 genes), the ROS/cytoskeletal organisation machinery (*NOX2, NOXR*), invasive growth regulators (*MIG1*), stress-response/conidiation regulator, polyketide synthases, and multiple genes from the Pmk1-pathway (*SLF1, RPP1, RPP5*) central to invasive growth (Cao et al., 2016, Lu et al., 2014, Mehrabi *et al*., 2008, Ryder *et al*., 2013, Osés-Ruiz et al., 2021). Bip1, however, uniquely activated a further 544 genes enriched for effectors (e.g. *SLP1*) and polyketide synthases (Mosquera *et al*., 2009, Takahashi *et al*., 2018), while Mst12 uniquely activated 347 genes involved in septation and polarity (Li *et al*., 2014, Liu et al., 2011, Mulford & Fassler, 2011, Shang *et al*., 2015, Jacob et al., 2014). These data support a model in which Bip1 drives an early metabolic and redox programme, Mst12 subsequently dominates appressorium maturation, and both factors converge to co-regulate the effector and invasive-growth machinery required for plant infection. Bip1 also has unique functions in controlling a distinct set of effectors and polyketide synthase genes, consistent with a role in orchestrating plant tissue colonisation (Fig. 5).

**Figure 5.**
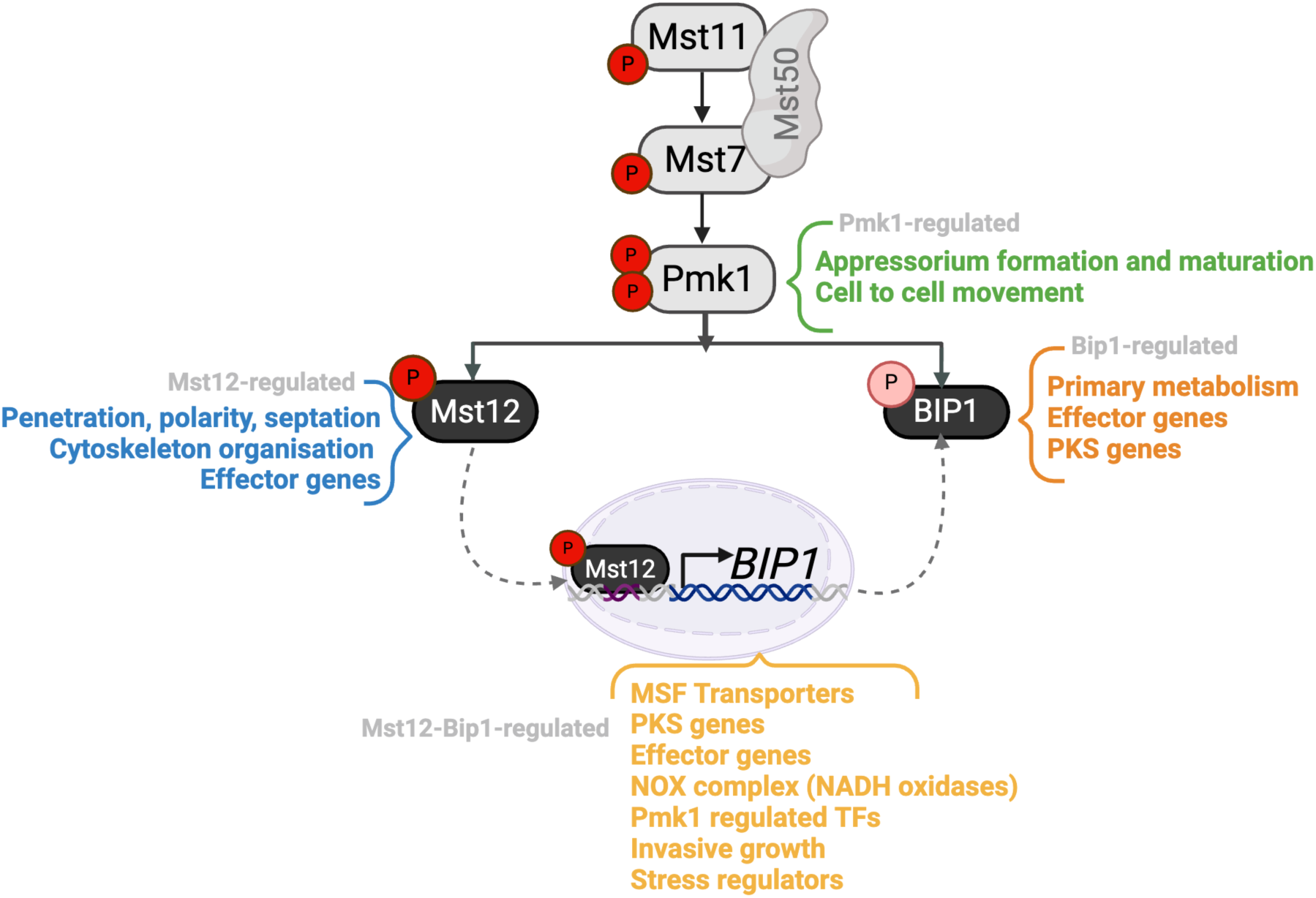
Model of a Pmk1-dependent Mst12-Bip1 transcriptional progam for rice blast infection. Three-layered MAPK activation, mediated by upstream sensors and a RAS-dependent phosphorelay, activates the Pmk1 MAPK, which is required for appressorium formation, maturation and invasive growth. Activated Pmk1 subsequently phosphorylates Mst12, which controls penetration, polarity, and septation through regulation of septin and actin ring assembly at the appressorium pore, while also repressing a number of effector proteins prior to plant infection. Mst12 binds a *cis*-regulatory element in the *BIP1* promoter to drive its transcription. Bip1 regulates primary metabolism during conidial germination its Mst12-mediated expression and Pmk1-dependent phosphorylation are required for effector regulation, PKS gene regulation and invasive growth. Together, Mst12 and Bip1 co-regulate a large set of infection-related genes, while Bip1 controls a distinct set of effectors associated with invasive growth by the rice blast fungus.

Repressed genes across all time-points meanwhile comprise transcription factors (Cao et al., 2016, Lu et al., 2014), polarity factors (Xu *et al*., 1998), proteins with DNA and RNA-binding domains, chromatin modifiers (*KMT6, HST1*) (Ding *et al*., 2010, Meng *et al*., 2021), and kinases (Yi *et al*., 2008), including Pmk1 itself, suggesting a potential negative feedback loop of the signalling cascade, as well as repression of genes associated with vegetative mycelial growth.

## Discussion

Fungi are unique among microbial pathogens in elaborating specialised infection structures called appressoria to breach the tough outer surface of plants (Ryder et al., 2022, Mendgen *et al*., 1996, Talbot, 2019). Extensive genetic and cell biological studies have identified numerous determinants of appressorium development in *M.oryzae* (Sakulkoo et al., 2018, Wilson & Talbot, 2009, Osés-Ruiz et al., 2021, Cruz-Mireles et al., 2024, Ryder et al., 2022), but how these individual components converge into the transcriptional programs that drive plant infection remains poorly understood.

We have shown that Bip1, a recently identified bZIP transcription factor (Lambou et al., 2024), plays a central role in both appressorium development and in preparing the fungus for biotrophic colonisation. bZIPs are a widely conserved family of transcription factors across eukaryotes (Vinson *et al*., 2006) with a characteristic DNA-binding basic region and a leucine zipper domain that mediates dimerization (McLachlan & Stewart, 1975). bZIP networks regulate development, stress responses and metabolism across pathogenic fungi (Son *et al*., 2011, Tian *et al*., 2011, Kong et al., 2015, Tang et al., 2015), and in *M. oryzae* have been linked to oxidative stress (e.g MoAPI), turgor regulation, effector expression, and virulence (Guo *et al*., 2011, Chen *et al*., 2014, Liu *et al*., 2022, Cao *et al*., 2022). Prior work, however, has relied on targeted assays such as RT-PCR and a single microarray analysis of the *Δbip1* mutant, which proposed that it acts independently of appressorium signal transduction pathways including the Pmk1 MAPK pathway (Lambou et al., 2024).

By contrast, combining phosphoproteomic, transcriptomic and cell biological experiments, we found evidence that Bip1 is an integral, Pmk1-dependent component of this network. A non-phosphorylatable version of Bip1 abolished virulence, demonstrating that Pmk1-dependent phosphorylation is essential for Bip1 function. Phosphorylation is the most common post-transcriptional modification for bZIP transcription factors, which can affect their homo or hetero-dimerization and subsequent activation/deactivation (Guo *et al*., 2024, Amoutzias *et al*., 2006, Schütze *et al*., 2008), and an analogous mechanism operates in *Aspergillus nidulans* where the bZIP transcription factor AtfA is phosphorylated by the p38/Hog1 MAPK pathway to regulate spore function (Lara-Rojas *et al*., 2011).

As the Mst12 regulator is a well-known substrate of Pmk1, we also examined its relationship to Bip1, despite it being reported to act independently (Lambou et al., 2024). Motif analysis, guided by published Mst12 ChIP-seq data (Osés-Ruiz et al., 2021), and Alphafold-3 modelling, identified a single Mst12-binding site in the *BIP1* promoter, predicted to interact through a C_2_H_2_ zinc finger region of Mst12, previously linked to its virulence function (Wong Sak Hoi & Dumas, 2010). EMSA confirmed Mst12 binding to the *BIP1* promoter element, and chimaeras lacking the motif were unable to restore pathogenicity when expressed in a *Δbip1* mutant, confirming that Mst12 acts as a direct upstream activator of *BIP1*. Our data therefore supports a model in which Pmk1 regulates Bip1 both by phosphorylation at a pathogen-conserved serine residue (S32) and through Mst12-dependent transcriptional activation of *BIP1*. The same Mst12-binding motif occurs in 658 other genes, including other bZIP transcription factors, that might also be transcriptionally-dependent on the Pmk1-Mst12 cascade.

How then could an Mst12-Bip1 transcriptional network operate to regulate plant infection? Few examples exist of bZIPs acting within transcription factor networks as complex as appressorium development, though comparable activation/repression regulation is found in other filamentous fungi. In *Cryptococcus neoformans*, for example, the Gsb1 bZIP protein can activate and repress a number of transcription factors required for virulence, mating and oxidative stress responses (Cheon *et al*., 2017) and, nitrogen assimilation in *A. nidulans* is regulated by the AreA/AreB complex, in which AreB acts with the MeaB bZIP TF to suppress AreA transcriptional activation (Michielse *et al*., 2014, John *et al*., 2021), while iron homeostasis regulator HapX competes with a GATA TF that suppresses iron assimilation (López-Berges *et al*., 2012, Jung *et al*., 2010). In *M. oryzae*, the Yap1-family bZIP MoAp1 regulates oxidative-stress and potentially cooperating with the response-regulator Skn7 (Mulford & Fassler, 2011, Rodrigues-Pousada *et al*., 2010), in a similar cooperative regulation as we describe for the Mst12-Bip1 network.

To investigate the precise relationship between Bip1 and Mst12, we carried out comparative global transcriptional profiling during appressorium development (0, 4 and 24h). This provided a temporal overview (Fig. 5) revealing that Bip1 dominates the earliest activation programme (metabolic and redox genes), Mst12 subsequently becomes the critical regulator of appressorium maturation and penetration machinery, and the two converge prior to host penetration, co-regulating a large set of genes including transcription factors, effectors, transporters and secondary-metabolite programmes required for plant colonisation (Cao et al., 2016, Lu et al., 2014, Osés-Ruiz et al., 2021, Ryder et al., 2013, Mehrabi et al., 2008). Many of the transcription factors co-regulated by Mst12 and Bip1 have not yet been functionally characterised, providing a platform of new regulator candidates of invasive growth. Bip1-specific regulation at this time shows a strong bias towards plant-tissue colonisation, with activation of the apoplastic effector *SLP1*, for example, which suppresses chitin-triggered immunity (Mentlak *et al*., 2012), as well as other previously characterised effectors (Mosquera et al., 2009, Takahashi et al., 2018), and polyketide synthase genes, consistent with secondary-metabolite induction associated with biotrophic invasion. By contrast, Mst12-specific gene expression is associated with regulation of septation and polarity required for penetration (Li et al., 2014, Liu et al., 2011, Mulford & Fassler, 2011, Shang et al., 2015, Jacob et al., 2014).

In summary, we show that Mst12 is the main regulator of appressorium maturation and penetration machinery, while Bip1 drives early metabolic and redox programmes and later biotrophy-associated genes. Bip1 and Mst12 together are necessary for the canonical appressorial invasion programme, Pmk1-dependent transcription factor expression, effector regulation, cell-wall remodelling, and secondary metabolite generation. Strikingly, 273 effector-encoding genes are regulated by Bip1/Mst12– constituting 50% of the total effector repertoire (Yan et al., 2023), of which 97 appear to be co-activated by both Bip1 and Mst12, 65 activated by Bip1 and just 35 activated solely by Mst12, demonstrating the predominant role of Bip1 and the shared Pmk1-Mst12-Bip1 cascade in deploying the *M. oryzae* effector repertoire. Together, these findings provide a model in which the Pmk1 MAPK pathway controls rice blast infection through a conserved transcriptional network centred on Mst12 and Bip1, linking MAPK signalling to gene expression programmes required for appressorium development, effector deployment and invasive growth of the blast fungus.

## Materials and methods

### Fungal strains, growth conditions, and preservation

*M. oryzae* isolates were maintained following established laboratory procedures, desiccated filter paper stocks stored at −20 °C were used to inoculate complete medium (CM) agar plates, and incubated at 24 °C under a 12 h light/12 h dark photoperiod (Molinari & Talbot, 2022, Talbot *et al*., 1993).

### Induction of appressorium development on glass coverslips

Conidia were harvested from 10-day-old CM cultures in 3 mL sterile water, filtered through Miracloth, and washed by centrifugation at 8,000 × g for 15 min. Pellets were resuspended and conidial concentration adjusted to 7.5 × 10⁴ conidia/mL with a haemocytometer (Improved Neubauer, UK) in the presence of 50 ng/µL 1,16-hexadecanediol (Sigma-Aldrich). For microscopy, 30 µL droplets were placed on hydrophobic borosilicate glass coverslips (Menzel-Gläser, Fisher Scientific UK) on water-soaked paper towels and incubated at 25 °C. For large-scale appressorium development assays, 8 hydrophobic coverslips were glued to square (120 mm) Petri filled with 50 mL of conidial suspension per time point and incubated at 25 °C without lids. Germinated samples were collected by scraping, snap-frozen in liquid nitrogen, and either processed immediately or stored at −80 °C; appressorium development was monitored using an inverted light microscope.

### Detached leaf pathogenicity assays

Pathogenicity was assessed by detached leaf-drop infection assays on barley cultivar Golden Promise. Conidia were harvested as described above and adjusted to 5 × 10⁴ conidia/mL in 0.2% (w/v) gelatin. Seven-day-old barley seedlings leaves were detached and placed on water-soaked Whatman (3mm) paper in square (120 mm) Petri dishes. Each leaf was inoculated with 3–4 droplets (20 µL each) of conidial suspension and incubated at 25 °C under a 12 h light/12 h dark photoperiod; disease symptoms were scored 5 days post-inoculation.

### Confocal laser scanning microscopy

Confocal imaging used a Leica TCS SP8 (63×/1.4 NA oil objective) with HyD detectors, using 488 nm excitation with emission collected at 500–550 nm for GFP and 520-530 nm for neonGreen, and 560 nm excitation with emission collected at 550–600 nm for dTomato. Images were acquired in Leica LAS AF software (Leica Microsystems, USA) and processed with Fiji/ImageJ2 version 2.14.0 (Rueden *et al*., 2017).

### Whole-genome sequencing and read processing

Genomic DNA was quantified using a NanoDrop (Fisher Scientific) and Qubit 2.0 (Thermo Fisher Scientific) and submitted to Novogene (Cambridge, UK) for whole-genome sequencing (350 bp insert, Illumina HiSeq 2500, paired-end). Raw reads were assessed with FastQC (Babraham Bioinformatics), trimmed with Trimmomatic (Bolger *et al*., 2014) and re-evaluated using FastQC. Reads were aligned to both a Guy11 and MG8 70-15 reference genome using BWA-MEM (bwa 0.7.7). SAM files were converted to BAM and sorted using SAMtools 1.9 (Li *et al*., 2009). Alignments were inspected in IGV (Robinson *et al*., 2011), against reference GFF3 annotations, querying genes by coordinate or MGG identifier.

### RNA sequencing and differential expression analysis

Total RNA (Qiagen RNAeasy Plant Mini Kit) from *M. oryzae* developmental stages was submitted to Novogene (Cambridge, UK) for bulk RNA-seq analysis (150 bp inserts, HiSeq 2500, paired-end). Transcript abundance was estimated with Kallisto (Bray et al., 2016), and differential expression with edgeR, incorporating Kallisto quantification uncertainty (Pimentel et al., 2017) using log₂ fold change ≥ 1 and adjusted p-value ≤ 0.05. Multidimensional scaling used the plotMDS function in limma, and heatmaps and Euler diagrams were generated with ggplot2 and eulerr (Wilkinson, 2012, Wilkinson, 2011). Pearson correlations were computed with corrplot (Hahsler *et al*., 2008), and GO term enrichment of differentially expressed genes was performed using the TopGO package (Alexa & Rahnenfuhrer, 2010).

### Motif analysis

Promoter motif occurrence was evaluated using FIMO server (MEME suite 5.5.7). Predicted promoter regions of *M. oryzae* genes were scanned using the built-in upstream *M. oryzae* MG8 dataset.

### Protein structure prediction and visualisation

*In silico* protein, protein–protein, and protein–DNA complex structures were predicted using AlphaFold3 (Abramson *et al*., 2024, Jumper *et al*., 2021) each with >=5 times seed replicates. The model with the highest mean pLDDT was selected for downstream analysis, with highly distorted regions cropped for clarity in representations. Structures were visualised and inspected using UCSF ChimeraX (Pettersen *et al*., 2021).

### Generation of plasmids for *M. oryzae* transformation

Gene coding regions were amplified from genomic DNA and cloned using In-fusion. *BIP1* (*MGG_08118*) with 1.5 kb of upstream promoter sequence was fused to C-terminus mNeonGreen in the linearised pscSUR vector (sulfonylurea resistance cassette) (Lindsay *et al*., 2016). Phospho-DEAD (S32-A32) and phospho-MIMETIC (S32-D32) alleles of *BIP1* alleles or the promoter motif delected or replaced by a poly-thymine sequence (*BIP1pSEQT*), were generated by site-directed primer design and multi-fragment In-Fusion cloning. Cytoskeletal markers (LifeAct, Sep5 and Chm1) and Histone marker (H1) were previously generated in the laboratory (Gupta et al., 2015).

### Bip1 knockout generation

Δ*bip1* mutants were generated in the Guy11 wild-type strain of *M. oryzae* by homologous recombination (Bird & Bradshaw, 1997, Weld *et al*., 2006). A 1.5kb 5’ flanking region of the *BIP1* ORF, followed by a *BASTA* resistance gene under a constitutive promoter, and then another 1.5kb 3’ flanking region of the *BIP1* gene was used for replacement of the original sequence through protoplasts transformation (Talbot et al., 1993). PCR and Sanger sequencing verified positive colonies. Two independent mutants, Δ*bip1_1* and Δ*bip1_2*, were confirmed through Illumina whole-genome sequencing and infection assays.

### Electrophoretic mobility shift assays

EMSAs assessed sequence-specific binding of recombinant Mst12 (generated with the TnT Quick Coupled Transcription/Translation system, Promega) to a double-stranded DNA probe containing three tandem copies of the Mst12 DNA-binding motif (CCTGTAAT) or a mutated motif (CAAATAAT) and synthesised with a 5′ 6-FAM label (Novogene). Oligonucleotides were annealed by heating to 95 °C for 5 min in nuclease-free water and cooling to room temperature overnight. Binding reactions (15 µL) contained 2 µL of labelled probe, 5–10 µL of Mst12 TnT reticulocyte lysate or no-lysate control, and nuclease-free water. Reactions were incubated for 30 min at room temperature, and complexes were resolved on native polyacrylamide gels (4–6% acrylamide:bisacrylamide, 29:1) in 0.5× TBE at 4 °C. Gels were pre-run for 20–30 min at 100 V, and after sample loading, electrophoresis was continued at 100 V for 30 min, fluorescent signals were detected using the Cy2 and Cy3 channels of an ImageQuant 800 imager (Cytiva).

### Network analysis

Network analyses were performed on differentially expressed genes from the RNA-seq comparisons explained above. DEGs were annotated using InterPro domains (Paysan-Lafosse *et al*., 2023) and known pathway annotations from *Cruz-Mireles et al., 2024* and assigned to broad functional categories (effectors, transcription factors etc.). The network layout was constructed using a custom script, which was imported to Gephi (Bastian *et al*., 2009), organised based on functional annotations using the ForceAltas2 layout algorithm (Jacomy *et al*., 2014). Network edges connected genes based on functional annotations and nodes were labelled as known gene names from MagnaGenes v.1.0 (Foster *et al*., 2021) where available. Genes with no annotated gene names were labelled with MGG identifiers.

## Acknowledgments

This work was supported by the Gatsby Charitable Foundation, a Biotechnology and Biological Sciences research council (BBSRC) doctoral training award BB/T008717/1 to CM and BBSRC Advancing Plant Health Institute Strategic Programme Grant to NJT BB/Y002997/1.

## Author Contributions

Author contributions: C.M and N.J.T. conducted the research and wrote the manuscript; C.M and N.S performed analysis of the transcriptomic data; L.R, X.Y, and V.G helped with experiments; C.M, N.S, L.R, X.Y, V.G, T.N and N.J.T contributed in editing the final version of the manuscript.

## Classification

BIOLOGICAL SCIENCES: Microbiology

Data deposition: https://github.com/nehasahu486/bip1_mst12_CM

