## Supplementary Figures for "A Pmk1-Regulated Mst12-Bip1 Network Coordinates Appressorium Function, Effector Deployment, and Invasive Growth by *Magnaporthe oryzae*"

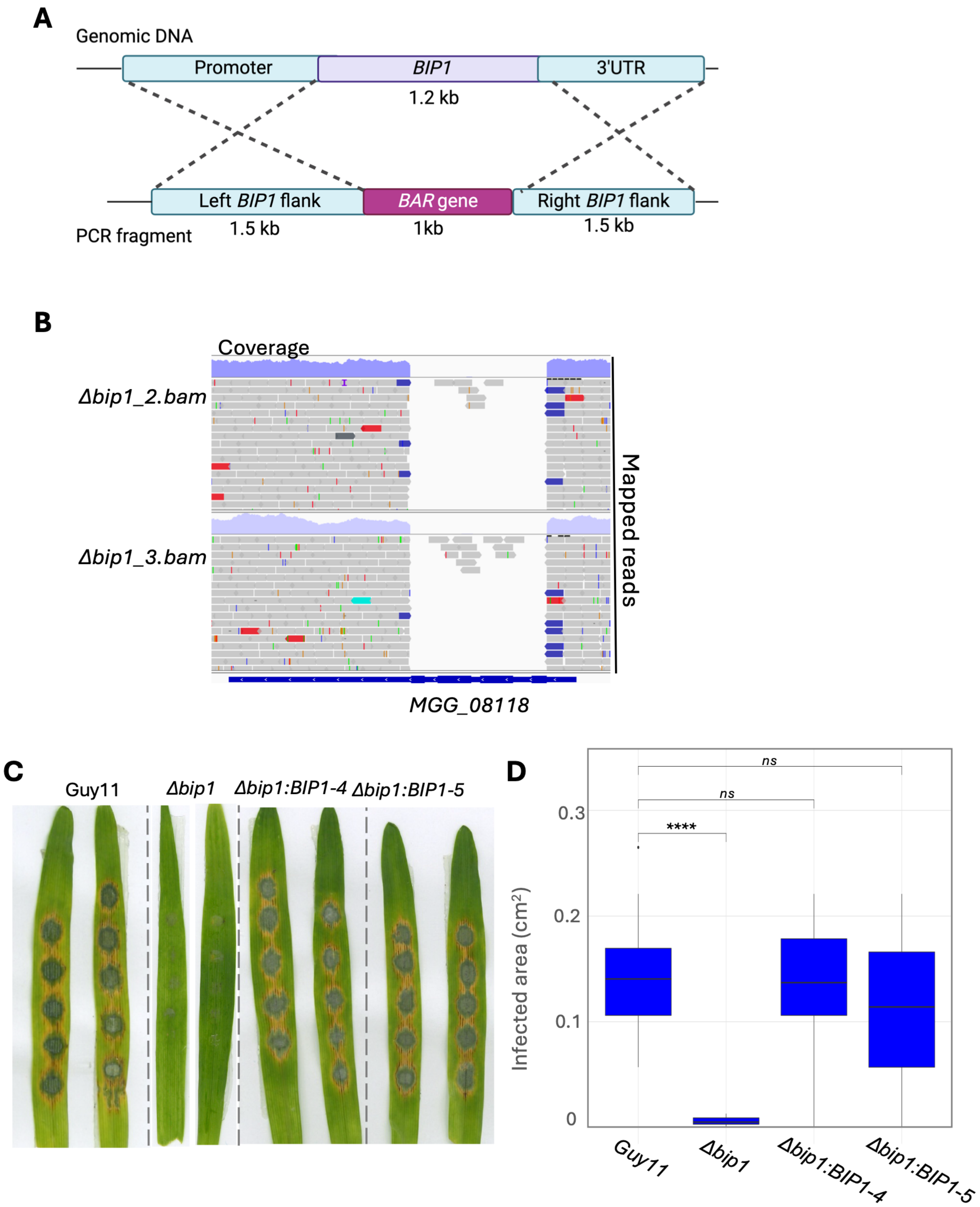


**Figure S1. Generation of *Δbip1 Magnaporthe oryzae* strains.** (A) DNA fragment designed to generate a *Δbip1* mutant through homologous recombination; 1.5kb 5' flanking region of the *BIP1* gene, followed by a BASTA resistance gene under a constitutive promoter, and then another 1.5kb 3' flanking region of the *BIP1* gene. (B) IGV snapshot of the *BIP1* (*MGG_08118*) coding region for two deletion mutants *Δbip1*_2 and *Δbip1*_3 confirms no read coverage at the *BIP1* coding region for these strains. (C) Detached leaf assays (barley cv. Golden Promise; see Methods) Guy11, *Δbip1,* and two independent complementation lines *Δbip1:BIP1-GFP_4* and *Δbip1:BIP1-GFP_5*., 5 dpi. (B) Box plots show the calculated disease area (n=20 lesions per replicate). The mean value is shown as a horizontal line inside the box. T-tests show significantly reduced virulence (p-value<0.05) for the *Δbip1* mutant but not for the complemented strains. Results represent four biological replicates.

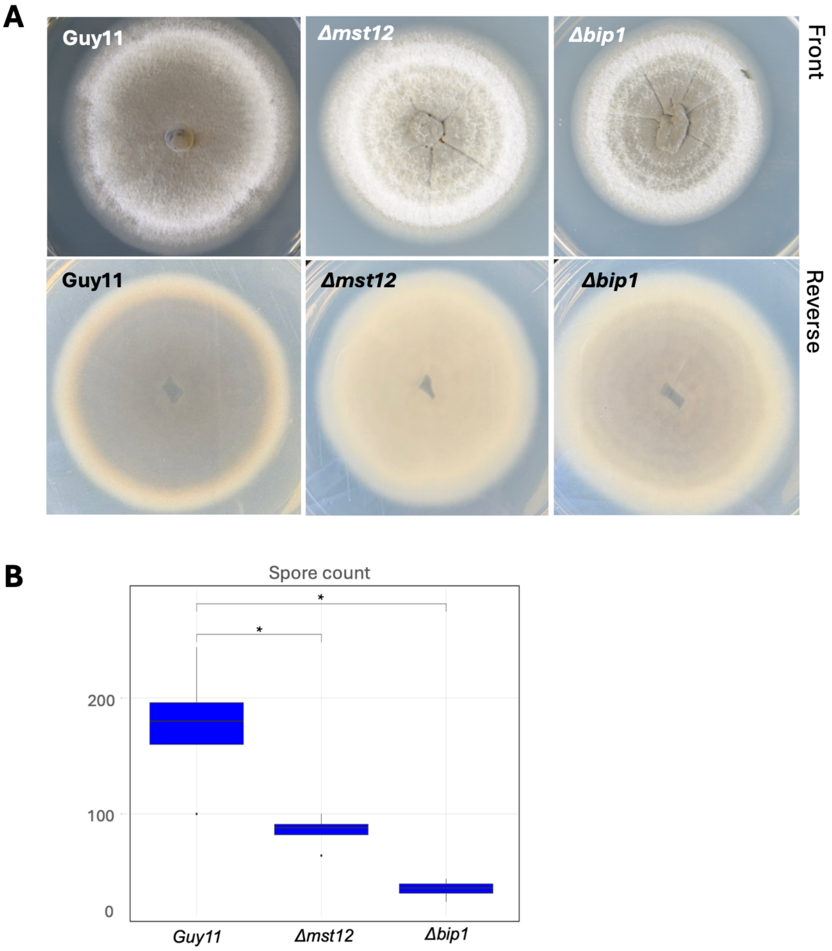


**Figure S2. *Δbip1* mutants show reduced growth rate and sporulation.** (A) Mycelial growth of Guy11, *Δmst12* and *Δbip1* on CM plates after 10 days post inoculation. Upper panel shows the front of the plate, and lower panel shows the reverse of the plate. (B) Box plots for the calculated spore count for each strain. The mean value is shown as a horizontal line inside the box. T-tests show significantly reduced sporulation (p-value<0.05) for both the *Δmst12 and Δbip1* mutants. Results represent three biological replicates.

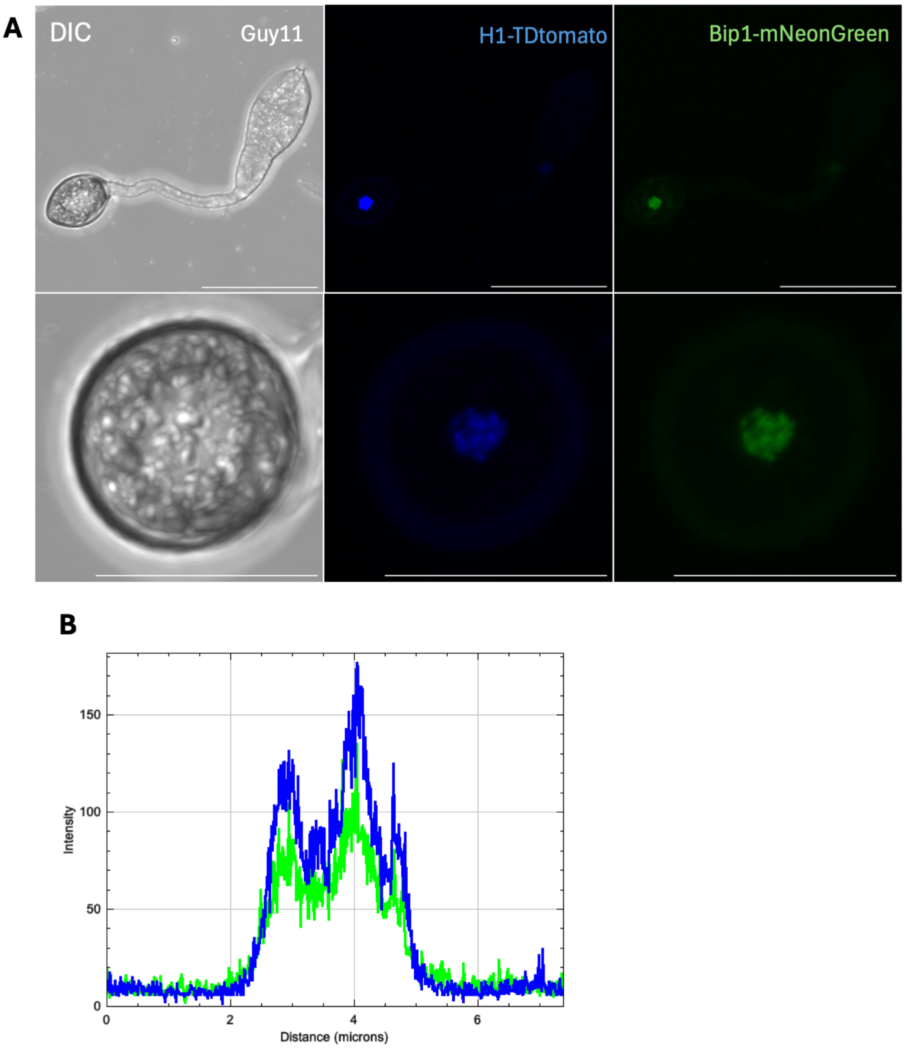


**Figure S3. Bip1 localises to the nucleus.** (A) Micrographs and (B) line scan of *M. oryzae* strain expressing histone H1-TDtomato and BIP1-mNeonGreen co-localising in the appressorium nucleus. Micrographs show a maximum projection of Z-stack images. Representative images from three biological replicates. Conidia were harvested, inoculated on hydrophobic coverslips, and observed by laser confocal microscopy at 24h. Scale bar = 20 μm.

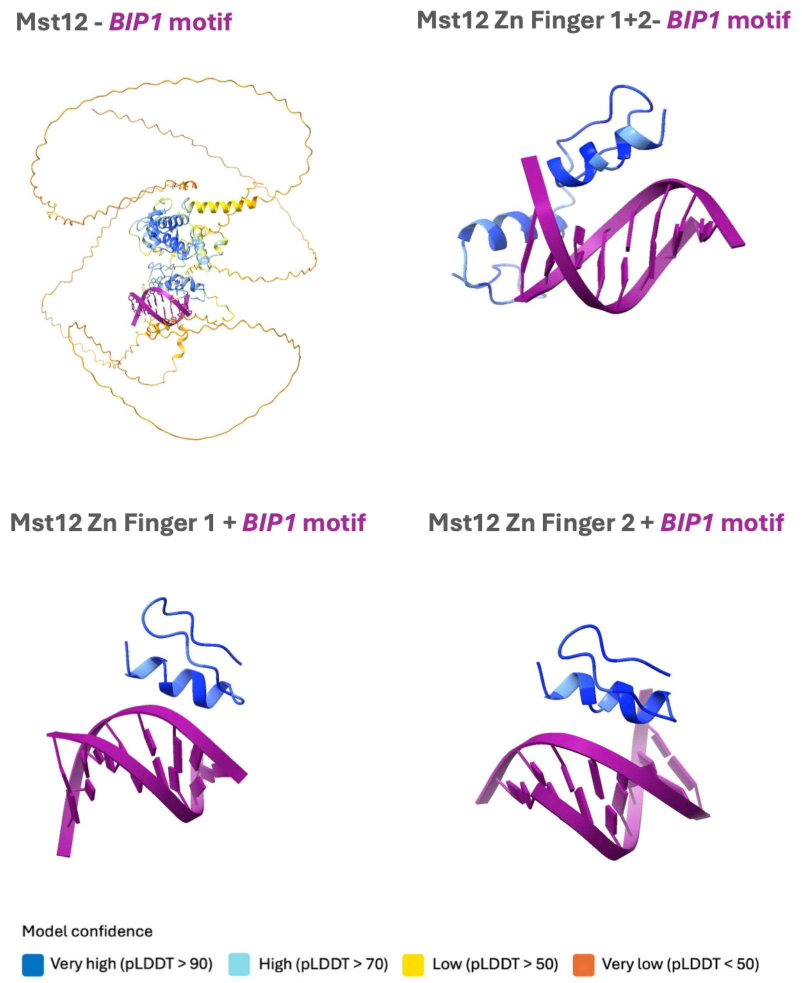


**Figure S4. Alphafold-3 predictions of Mst12 and the *BIP1* motif.** Four panels showing the best-ranked model predicted through Alphafold-3 of the Mst12 in complex with the *BIP1* promoter motif: full-length Mst12, the $C_2H_2$ zinc finger domain, the first $C_2H_2$ zinc finger, and the second $C_2H_2$ zinc finger. pLDDT scores are shown with the model confidence colouring. Terminal distorted regions have been cropped for visualisation purposes.

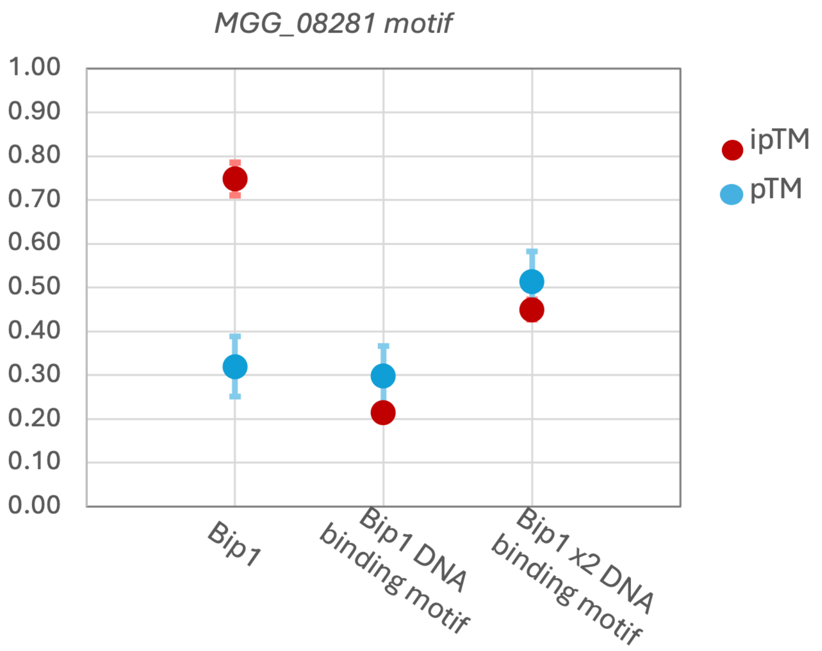


**Figure S5. Alphafold-3 predictions of Bip1 and its cognate DNA-interacting motif found in *MGG_08281*.** A scatter plot shows pTM and ipTM values for three AlphaFold 3 predictions: full-length Bip1 with one Bip1-binding motif from the *MGG_08281* promoter, the DNA binding motif with a Bip1 chimaera (with only the DNA binding region of the protein) and the DNA binding motif with two Bip1 chimaeras. All predictions were repeated 5 times with different seed values.

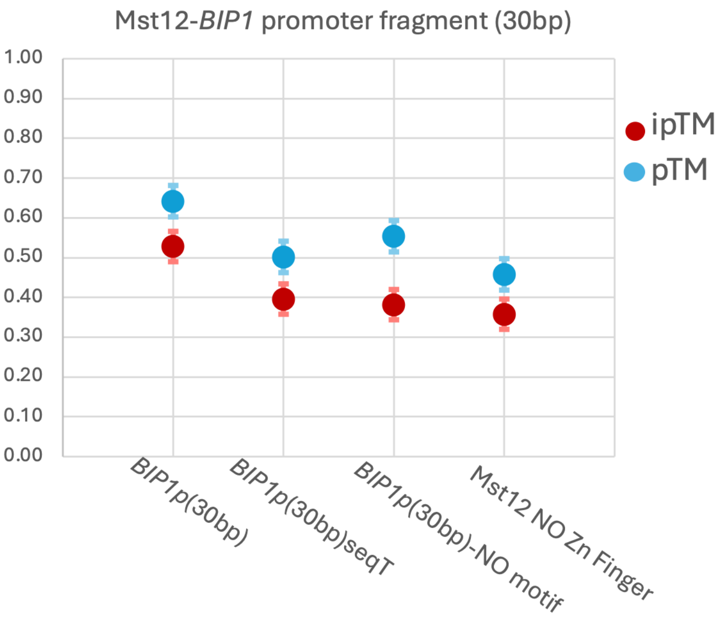


**Figure S6. AF3 predictions of Mst12 and a fragment (30bp) of the *BIP1* promoter containing the Mst12-DNA binding motif.** A scatter plot shows pTM and ipTM values for four AlphaFold 3 predictions of Mst12 with a 30 bp *BIP1* promoter fragment containing the Mst12-binding motif, the same 30 bp promoter fragment lacking the motif, the promoter fragment in which the motif is replaced by a poly-T sequence, and a chimeric Mst12 lacking the Zn-finger domain with the 30 bp BIP1 promoter fragment. All predictions were repeated 5 times with different seed values.

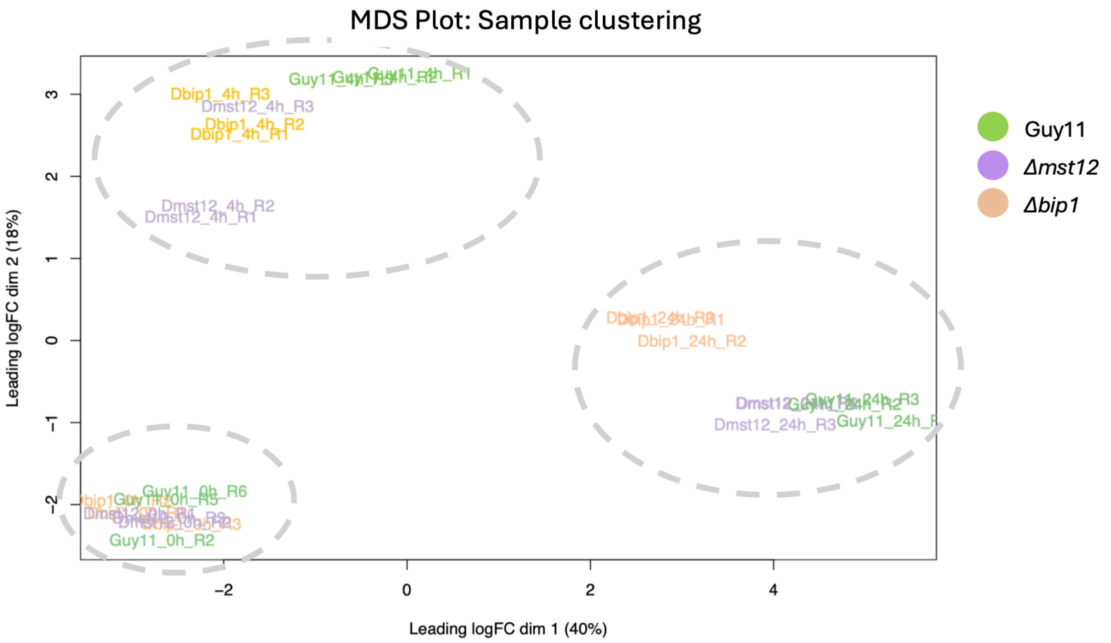


**Figure S7. Multidimensional scaling (MDS) plot for the time-course RNA-Seq analysis.** MDS plot was generated to analyse sample clustering based on expression value. Combining both dimensions, there is a 58% variance across samples. Biological replicates, represented by the same colours, cluster together, suggesting reproducibility of the data. Grouping is also observed by time point (0h, 4h, 24h) of appressorium development. Guy11 samples are represented in green, *Δmst12* samples are represented in purple, and *Δbip1* samples are represented in orange.

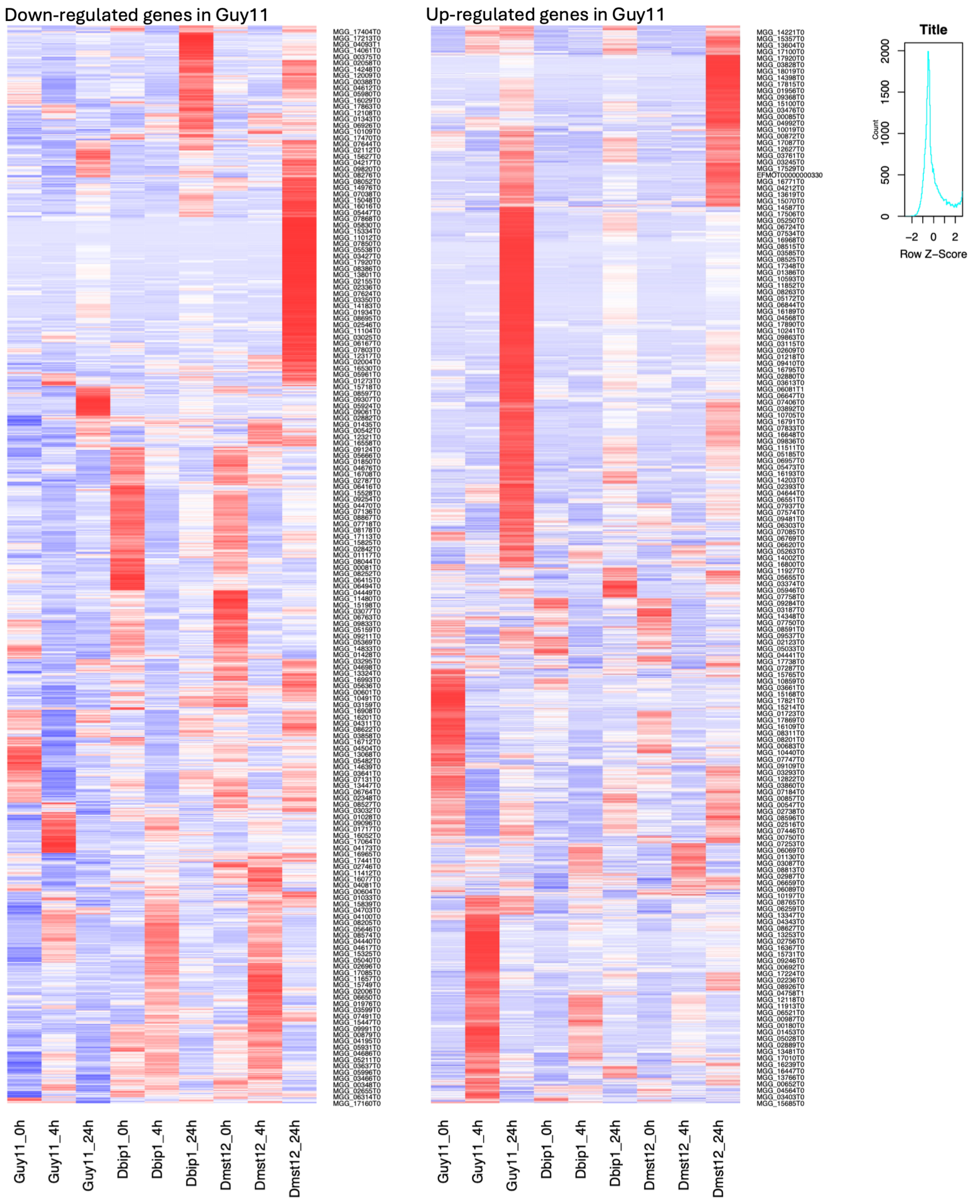


**Figure S8. Heatmap of DEGs.** Heatmap of down-regulated (left) and up-regulated (right) genes in Guy11 identified from global transcriptomic analysis of *Δmst12* and *Δbip1* mutants during appressorium development. Row Z score = (TPM for a single gene) – mean/standard deviation.

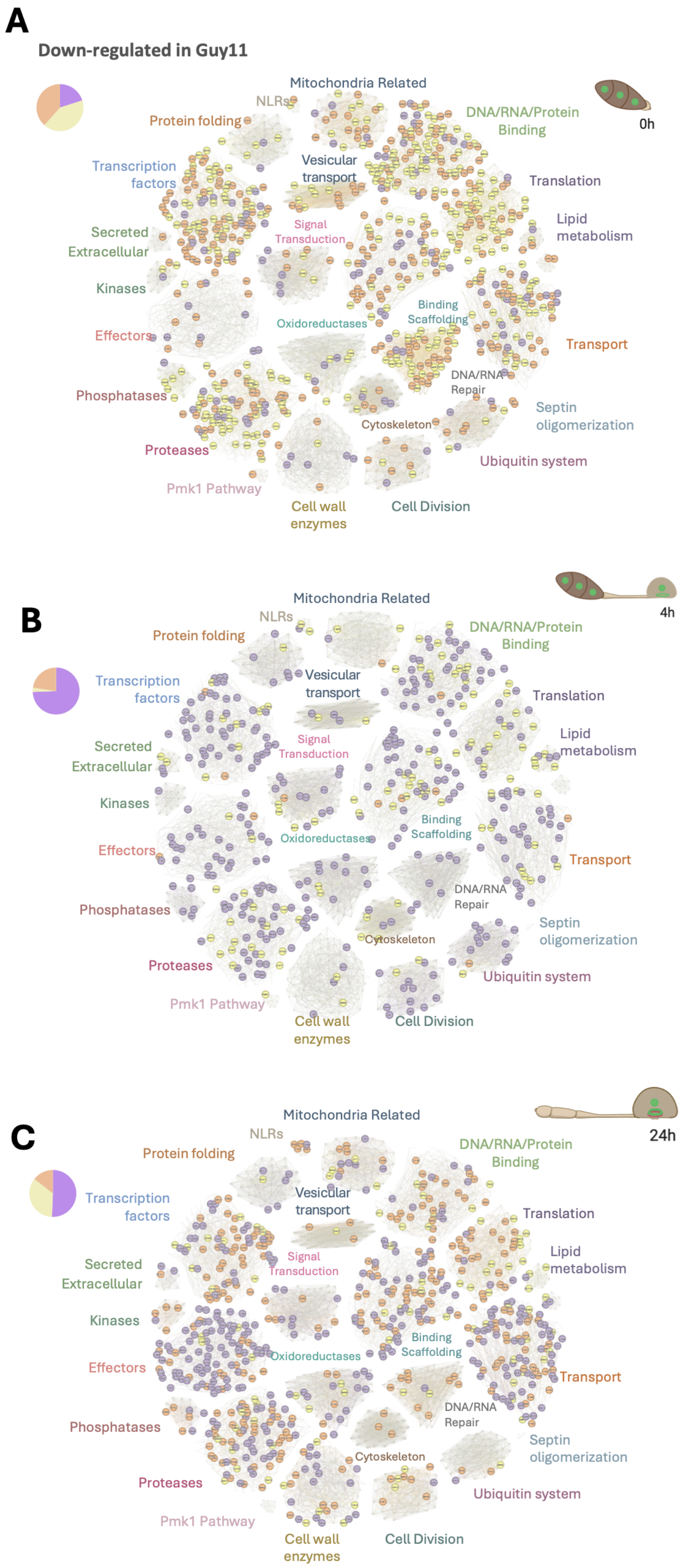


**Figure S9. Network analysis of downregulated-DEGs.** The base network shown in Figure 4D was used to plot genes down-regulated in Guy11 for the *Δbip1* mutant (orange), *Δmst12* mutant (purple), and both *Δbip1* and *Δmst12* mutants (yellow) at 0 h (A), 4 h (B), and 24 h (C) of appressorium development.
